# Claudin-5 binder enhances focused ultrasound-mediated opening in an *in vitro* blood-brain barrier model

**DOI:** 10.1101/2021.08.01.454692

**Authors:** Liyu Chen, Ratneswary Sutharsan, Jonathan LF Lee, Esteban Cruz, Blaise Asnicar, Tishila Palliyaguru, Gerhard Leinenga, Jürgen Götz

## Abstract

**Rationale:** The blood-brain barrier (BBB) while functioning as a gatekeeper of the brain, impedes cerebral drug delivery. An emerging technology to overcome this limitation is focused ultrasound (FUS). When FUS interacts with intravenously injected microbubbles (FUS^+MB^), the BBB opens, transiently allowing the access of therapeutic agents into the brain. However, the ultrasound parameters need to be tightly tuned: when the acoustic pressure is too low there is no opening, and when it is too high, bleeds can occur. We therefore asked whether BBB permeability can be increased by combining FUS^+MB^ with a second modality such that in a clinical setting lower acoustic pressures could be potentially used.

**Methods:** Given that FUS achieves BBB opening by the disruption of tight junction (TJ) proteins such as claudin-5 of brain endothelial cells, we generated a stable MDCK II cell line (eGFP-hCldn5-MDCK II) that expresses fluorescently tagged human claudin-5. Two claudin-5 binders, mC5C2 (a peptide) and cCPEm (a truncated form of an enterotoxin), that have been reported previously to weaken the barrier, were synthesized and assessed for their abilities to enhance the permeability of cellular monolayers. We then performed a comparative analysis of single and combination treatments.

**Results:** We successfully generated a novel cell line that formed functional monolayers as validated by an increased transendothelial electrical resistance (TEER) reading and a low (< 0.2%) permeability to sodium fluorescein (376 Da). We found that the binders exerted a time- and concentration-dependent effect on BBB opening when incubated over an extended period, whereas FUS^+MB^ caused a rapid barrier opening followed by recovery after 12 hours within the tested pressure range. Importantly, preincubation with cCPEm prior to FUS^+MB^ treatment resulted in greater barrier opening compared to either FUS^+MB^ or cCPEm alone as measured by reduced TEER values and an increased permeability to fluorescently labelled 40 kDa dextran (FD40).

**Conclusion:** The data suggest that pre-incubation with clinically suitable binders to TJ proteins may be a general strategy to facilitate safer and more effective ultrasound-mediated BBB opening in cellular and animal systems and potentially also for the treatment of human diseases of the brain.

## Introduction

One of the biggest challenges in developing therapeutics for central nervous system (CNS) disorders is to achieve sufficient blood-brain barrier (BBB) penetration [1]. It has been estimated that the BBB presents an obstacle for more than 98% of small-molecule drugs and nearly all large-molecule therapeutics for CNS disorders [2]. The use of low-intensity focused ultrasound (FUS) in combination with intravenously injected microbubbles (FUS^+MB^) has been shown to be a targeted, non-invasive technique that can transiently disrupt the BBB to allow access of therapeutics to the brain [3, 4]. However, for safe and effective opening, the ultrasound parameters need to be tightly tuned. When the acoustic pressure is too low there is no opening, and when the pressure is too high this can potentially cause tissue damage such as microhaemorrhages and oedemas due to inertial cavitation of the microbubbles [5, 6]. We therefore investigated, using a novel *in vitro* BBB model, whether permeability can be increased by combining FUS with a second modality such that in a clinical setting lower acoustic pressures could be potentially used.

At the cellular level, the BBB is represented by the neurovascular unit (NVU) that includes endothelial cells, pericytes and astrocytes, as well as an ensheathing basement membrane. The NVU generates a dynamic interface between the circulation and brain parenchyma, providing anatomical and physiological protection for homeostasis of the CNS [1, 3, 7]. Transport across the brain endothelium is tightly controlled via (i) a paracellular barrier represented by interendothelial tight junctions (TJs); (ii) a transcytoplasmic barrier, owing to the unique properties of the brain capillary endothelial cells (BCECs) with a low level of endo- and transcytosis; (iii) an enzymatic barrier; and (iv) active multidrug efflux transporters. FUS^+MB^ achieves transient BBB opening both by facilitating transcytoplasmic transport and by severing the homophilic interactions between TJ proteins.

The TJs of BCECs are formed by a group of proteins which includes occludin, ZO-1 and the claudin family, of which claudin-5 is the most highly expressed [8] **(Fig. 1A)**. Claudin-5 has four transmembrane domains, a short cytoplasmic amino-terminus, two extracellular loops (ECL1 and ECL2), an intracellular loop and a longer cytoplasmic carboxy-terminus **(Fig. 1A)**. The ECLs contribute to homo- and heterophilic interactions, thereby tightening the paracellular space [9, 10].

**Figure 1.**
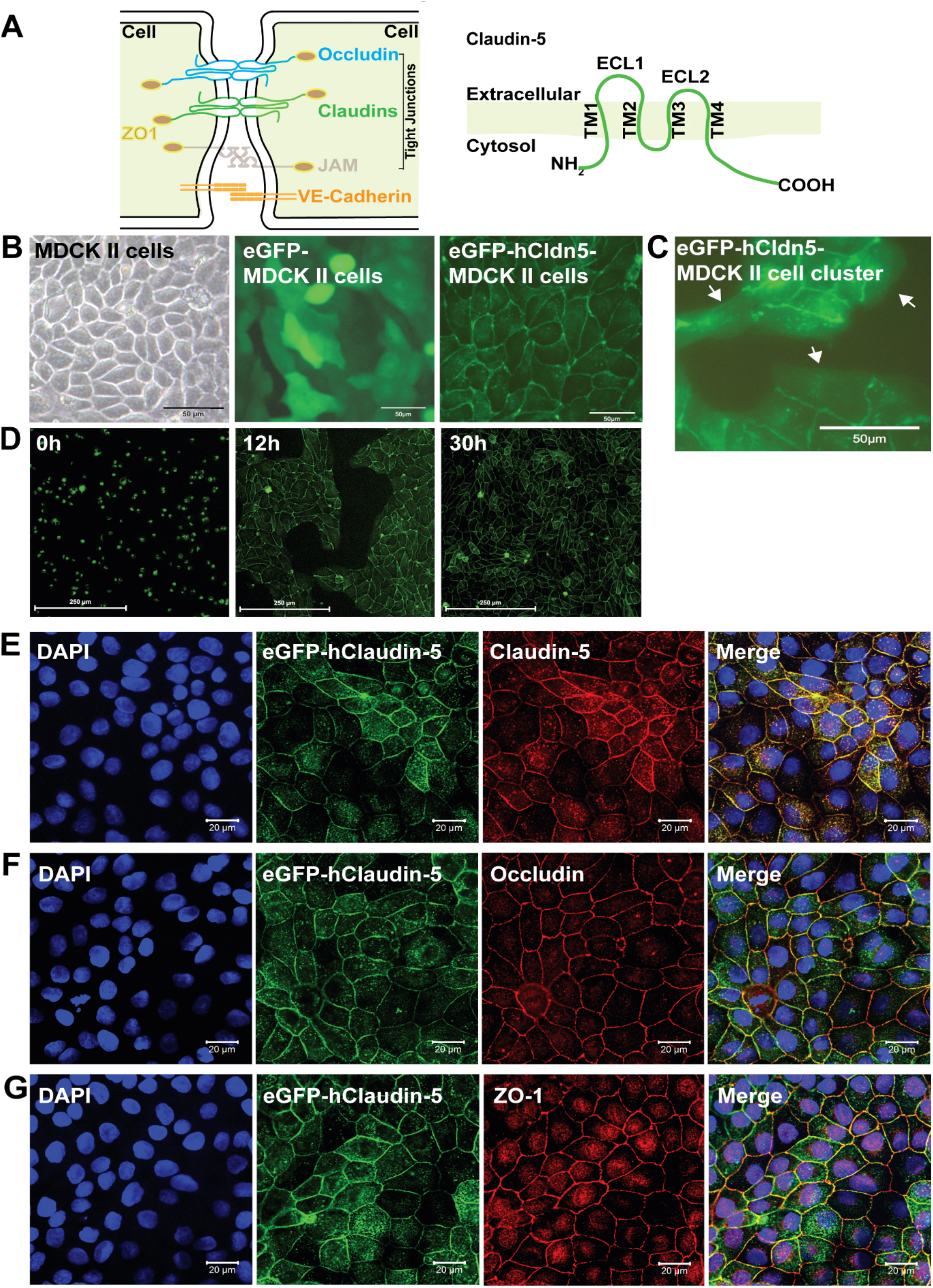
eGFP-hCldn5-MDCK II cells exhibit a tight monolayer and human claudin-5 is localized to cell/cell contacts. **(A)** Scheme of tight junctions formed by proteins such as claudin-5, occludin, and ZO-1. Domain structure of claudin-5. **(B)** Representative phase contrast image of confluent parental MDCK II cells and epifluorescence images of confluent MDCK II cells expressing eGFP only (eGFP-MDCK II) or eGFP-tagged human claudin-5 (eGFP-hCldn5-MDCK II). **(C)** Epifluorescence images of isolated clusters of eGFP-hCldn5-MDCK II cells. White arrows indicate the absence of localization of eGFP fluorescence in areas without cell/cell contact. **(D)** Time-lapse fluorescence images of eGFP-hCldn5-MDCK II cells from the time of seeding at a density of 200,000 cells/cm^2^ (0 h) to complete formation of a confluent monolayer (30 h). **(E)** Expression of claudin-5, **(F)** occludin and **(G)** ZO-1, localized to cell/cell borders. Nuclei were stained with DAPI. Scale bars: 50 μm (A, B), 250 μm (C) and 20 μm (D-F).

Several *in vitro* models which recapitulate critical physiological parameters and molecular aspects of the NVU are available for studying the BBB. Analysis tools include immunolabelling of TJ proteins, and when cultured in Transwell inserts, allows measurement of the transendothelial electrical resistance (TEER) and leakage of cargoes [11]. TEER values are strong indicators of the integrity of cellular barriers before they are evaluated for transport of drugs or chemicals [12]. The TEER values measured *in vivo* have been reported to be as high as 5,900 Ω·cm^2^ [13]. In a co-culture system with human induced pluripotent stem cell (hiPSC)-derived endothelial cells (iBECs) and neuronal progenitor cells treated with retinoic acid, TEER values up to 5,000 Ω·cm^2^ have been achieved [14]. However, having an ongoing supply of iBECs is both costly and challenging [15]. On the other hand, cell lines such as immortalized human cerebral microvascular endothelial cells (hCMEC/D.3), co-cultured with astrocytes, yield a relatively low TEER value of 140 Ω·cm^2^ [16]. Here, we generated a Madin-Darby canine kidney (MDCK II) cell line that stably expresses fluorescently tagged human claudin-5 and has around three-fold higher TEER values than the parental cell-line, to assess BBB weakening and cargo leakage in response to different interventions.

Given the central role of claudin-5 in the NVU, several peptide-based approaches have been previously developed to target claudin-5 and thereby weaken the BBB. Dithmer and colleagues designed peptides based on ECL1 [17]. Of these, peptide mC5C2 displayed a nanomolar affinity for claudin-5 and achieved a size-selective (up to 40 kDa dextran) and reversible (12–48 h) increase in paracellular transport of cargoes across brain endothelial and claudin-5-transfected epithelial cell monolayers. Furthermore, the peptide safely opened the murine BBB *in vivo*, as demonstrated by leakage of a contrast agent using magnetic resonance imaging.

Different from mC5C2, the non-toxic carboxy-terminal domain of the clostridium perfringens enterotoxin, cCPE, binds to ECL2 of the subtype-specific claudins, claudin-3 and claudin-4, with high affinity, increasing paracellular permeability and enhancing drug absorption [18, 19]. Because cCPE does not bind to claudin-5, Protze and colleagues performed site-directed mutagenesis and generated a novel cCPE mutant (cCPE_Y306W/S313H_, cCPEm) with nanomolar affinity, which achieved transient BBB opening by selectively binding to claudin-5 [20]. cCPEm decreased TEER in a concentration-dependent and reversible manner in three *in vitro* BBB models derived from three species [21].

Here, we investigated the effects of FUS^+MB^ and the two claudin-5 binders, mC5C2 and cCPEm, both alone and in combination, on BBB properties in an MDCK II cell culture model we had generated that expresses fluorescently tagged human claudin-5. Our results revealed that pre-incubation with cCPEm improves the degree of opening for a range of acoustic pressures below 0.3 MPa (required for barrier opening) as determined by TEER and leakage of fluorescently labelled cargoes.

## Materials and methods

### Claudin-5 peptidomimetics and generation of the GST-cCPEm fusion protein

The claudin-5 peptidomimetic mC5C2 corresponding to residues 53-81 of ECL1 of murine claudin-5 was synthesized by GeneScript, using standard solid-phase peptide synthesis and fluorenylmehyloxycarbonyl (Fmoc) chemistry on a preloaded Fmoc-amino acid resin [17]. The peptide was carboxy-terminally amidated and had the expected mass and a purity >95%.

The GST-cCPEm fusion protein (**Suppl. Figure 1**) was generated by subcloning the carboxy-terminal domain of CPE (aa 194-319 with two mutations Y306W and S313H) via the *EcoRI* and *SalI* restriction sites of the pGEX-4T1 (GE Healthcare, cat. #28-945-9545-49) plasmid, in-frame with an amino-terminal GST tag to facilitate protein purification, thereby generating plasmid pGEX-4T1-cCPEm.

Next, BL21(DE3) *E. coli* (New England Biolabs, cat. #C25271) were transformed with pGEX-4T1-cCPEm and grown to OD600 ~ 0.6, with expression being induced by adding isopropy1-β-D-thiogalactopyranoside (IPTG, Meridian Bioscience, cat. #BIO-37036) to a final concentration of 1 mM. After 6 h at 30 °C with constant vigorous shaking, the bacteria were harvested by centrifugation (6,000 g, 20 min, 4 °C) and stored at −80 °C until purification. The bacterial pellet was resuspended in lysis buffer (25 mM tris-HCl, pH 8; 150 mM NaCl, 5 μM DTT) containing protease inhibitor (Complete Mini, EDTA-free; Roche Applied Science), 10 U/ml benzonase Nuclease (Sigma, cat. #E1014) and 100 μg/ml lysozyme (AMRESCO, cat. #0663-10G), and incubated on ice for 20 min. Cells were then lysed by intermittent sonication (Sonics Vibra-Cell VCX130) at a 60% amplitude for 3 min. The cell lysates were centrifuged at 20,000 g for 30 min at 4 °C and the supernatant was filtered through a 0.22 μm syringe filter (Millipore).

Purification of the GST-tagged recombinant protein was performed as described previously [22, 23]. Briefly, the supernatant of the bacterial lysate was purified by affinity chromatography using an automated Profinia Protein Purification system (Bio-Rad), followed by size-exclusion chromatography using a Superdex 200 Increase 10/300 GL column (GE Healthcare, cat. #28990944) with an ÄKTApurifier chromatography system (GE Healthcare) in 1×PBS with 1 mM DTT. A_280_ peak fractions were collected and assayed for GST expression by Coomassie staining and immunoblotting with an anti-GST antibody. GST-cCPEm containing fractions were pooled. The protein concentration was determined using a NanoDrop 2000 spectrophotometer (Thermo-Fisher) using a molar extinction coefficient of 73,185 M^−1^cm^−1^, as calculated by the online Expasy ProtParam portal. Aliquots of GST-cCPEm were stored at −80 °C until use.

### Molecular cloning, lentiviral particle production and generation of eGFP-hCldn5-MDCK II cells

To generate an MDCK II cell line that expresses eGFP-tagged human claudin-5 (hCldn5), a lentiviral vector, pLV-eGFP-hCldn5, was generated. First, a gBlock was generated for hCldn5, which included an amino-terminal glycine/serine-rich linker sequence. The gBlock was then amplified by PCR to introduce *KpnI and MluI* restriction sites. Next, eGFP was amplified from plasmid pLV-eGFP (Addgene cat. #36083) and the flanking *NheI* and *KpnI* restriction sites were introduced. The amplified products were digested and ligated with T4 DNA ligase (NEB cat. #B0202S) into pLX_311-KRAB-dCas9 (Addgene cat. #96918) using the *NheI* and *MluI* restriction enzymes to generate pLX-eGFP-hCldn5, followed by sequencing for verification purposes. As a negative control, we constructed an empty eGFP lentiviral vector (pLX-eGFP).

The experimental procedure to generate active lentiviral particles has been published previously [24]. Briefly, adherent Lenti-X cells (Takara Bio, #632180) were cultured in Dulbecco’s Modified Eagle Medium (DMEM) with pyruvate containing 10% foetal bovine serum (FBS). The third generation lentiviral packaging system plasmids pMDLg/pRRE (Addgene, cat. #12251), pRSV-Rev (Addgene, cat. #12253) and envelope expressing plasmid pMD2.g (Addgene, cat. #12259) were used to transfect Lenti-X cells by CaPO_4_ precipitation [25, 26]. The lentivirus-containing medium was collected after 48 h and 72 h, centrifuged at 3,000 g for 5 min and filtered at 0.45 μm. For the final purification and concentration, the lentivirus-containing medium was suspended above a 10% sucrose cushion (100 mM NaCl, 0.5 mM EDTA, 10% sucrose, 50 mM Tris-HCl to pH 7.4) and centrifuged at 10,000 g for 4 h at 4 °C. The supernatant was discarded, the lentiviral pellets were resuspended in 100 μl of 1× HBSS (Hank’s balanced salt solution, Thermo-Fisher, cat. #14175095), and 20 μl aliquots were snap-frozen in liquid nitrogen and then stored at −80 °C until use. As a negative control, lentiviral particles packaged with pLX-eGFP were generated.

To generate a stable eGFP-hCldn5-MDCK II cell line, MDCK II cells were transduced with active lentiviral particles for 72 h. Infected cells were then exposed to growth medium containing 8 μg/ml blasticidin (Sigma, cat. #15205) for up to 14 days (**Suppl. Figure 2)**.

### Cell culture and additional cell-lines

MDCK II cells were purchased from the European Collection of Authenticated Cell Culture (ECACC, cat. #00062107) and used to generate a cell line that stably expresses human claudin-5 amino-terminally tagged with eGFP. All MDCK II cell lines in this study were cultured in Minimum Essential Medium Eagle (MEM) (Sigma, cat. #M4655) supplemented with 5% FBS. Lenti-X 293T cells were used for lentiviral particles production and cultured in DMEM or DMEM supplemented with 1mM pyruvate (Thermo-Fisher, cat. #11995073) and 10% FBS. All culture media contained 100 U/ml penicillin and 100 U/ml streptomycin (Thermo-Fisher, cat. #15070063). Dulbecco’s phosphate-buffered saline (DPBS, Thermo-Fisher, cat. #14190144) was used for washing steps. All cell lines were maintained in 5% CO2 at 37 °C.

iBECs were derived from hiPSCs. hiPSCs, a generous gift provided by Dr. Anthony White (QIMR Berghofer Medical Research Institute, Australia) were maintained on human recombinant vitronectin in StemFlex medium (Life Technologies, cat. # A3349401). Differentiation was performed as previously described [15, 27]. Briefly, cells were plated on Matrigel (In Vitro Technologies, cat. # FAL354277) coated plates in StemFlex medium supplemented with 10 μM ROCK inhibitor (StemCell Technologies, cat. #. 72304). Three days after plating, the culture medium was changed for unconditioned medium consisting of DMEM/F12+GlutaMAX, 20% Knockout Serum Replacement (Life Technologies, cat. #.10828028), 1× non-essential amino acids (Sigma, cat. #M7145) and 0.1 mM β-mercaptoethanol (Sigma, cat. #M3148-25ml) to induce spontaneous differentiation. After 6-8 days in unconditioned medium, the culture medium was changed to endothelial cell medium (ECM) supplemented with 2% B27 (Life Technologies, cat. #17504-044), 20 ng/ml basic fibroblast growth factor (bFGF) (Peprotech, cat. #100-18B-250) and 10 μM retinoic acid (Sigma, cat. #R2625). Cells were maintained in supplemented ECM for 2-3 days, after which they were transferred onto collagen IV (Sigma, cat. #C5533) and fibronectin (Life Technologies, cat. #33016015) coated Transwell inserts (Corning Inc.). After a further 2 days, the cells were maintained in ECM+B27 medium without bFGF and retinoic acid for one additional day. TEER reading was performed on these iBECs 48 h after subculturing, with cells displaying 100% confluency.

### Validation of GST-cCPEm binding of claudin-5 in isolated cerebral microvessels

To validate GST-cCPEm binding to endogenous claudin-5, cerebral microvessels were isolated from C57Bl6/J mice, with modification of a previously published protocol [28] **(Suppl. Fig. 3A)**. Briefly, animals were euthanized by intraperitoneal injection of 350 mg/kg pentobarbitone (Lethabarb, Virbac), after which their brains were dissected and submerged in MCDB131 medium (Thermo-Fisher, cat. #10372019). Meninges and meningeal vessels were removed by gently rolling the brains on damp blotting paper (Thermo-Fisher, cat. #14190144). The grey matter was dissected out using a razor blade and curved forceps, followed by Dounce homogenization in MCDB131 medium using a tissue grinder. The suspension was centrifuged at 2,000 g for 5 min at 4 °C (BeckmanCoulter, Avanti J-26 XPI) to remove myelin debris. The pellet was then resuspended in Dulbecco’s phosphate buffered saline (DPBS) containing 25% w/v 70 kDa dextran, and then centrifuged at 10,000 g for 15 min at 4 °C to sediment the red microvessel pellet. The step was repeated for further white matter removal. The pellets were resuspended in MCDB131, transferred to 40 μm cell strainers, and washed with DPBS. The filters were then inverted and microvessels were retrieved with MCDB131 solution containing 0.5% bovine serum albumin (BSA). Cellular composition and purity were validated for expression of glial fibrillary acidic protein (GFAP) and platelet derived growth factor receptor beta (PDGFRβ) to confirm the presence of astrocytes and pericytes, respectively, and of class III beta-tubulin (TUJ1) to confirm the absence of neurons **(Suppl. Fig. 3B-D)**.

To determine GST-mCPE binding, the purified microvessels were plated onto Superfrost Plus microscope slides (Menzel-Gläser, cat. #SF41296SP) and left to air-dry at room temperature for 30 min. They were then incubated with 100 μg/ml GST-cCPEm diluted in MCDB131 + 0.5% BSA in a 37 °C incubator for 30 min, followed by three washes with DPBS containing 0.1% NP-40. They were then fixed with 4% PFA-DPBS for 15 min, permeabilised with DPBS + 0.1% NP-40 for 15 min and blocked with 5% BSA-DPBS at room temperature for 1 h. The microvessels were subsequently incubated with an anti-mouse GST primary antibody at 4 °C overnight, washed three times with DPBS + 0.1% NP-40, stained with a goat-anti-mouse Alexa Fluor 594 secondary antibody for 1 h at room temperature, and counterstained with 4′,6-diamidino-2-phenylindole (DAPI). Slides were mounted with Vectashield antifade mounting medium and images captured using a Plan Apochromat 63×/1.4 NA oil-immersion objective on a Zeiss LSM 710 confocal microscope built around a Zeiss Axio Observer Z1.

### Dextran and sodium fluorescein cargoes

Fluorescein isothiocyanate-dextrans (FDs) and sodium fluorescein (NaFl) were used for permeability studies. FD4 has a molecular weight of 4 kDa (Sigma cat. #FD4-100MG) with a hydrodynamic radius (R_H_) ~0.95 nm, and FD40 has a molecular weight of 40 kDa (Thermo-Fisher, cat. #D1845) with a hydrodynamic radius (R_H_) ~6 nm. The dextrans were diluted to a stock concentration of 10 mg/ml in PBS. NaFl (Sigma, cat. #F6377) with a molecular weight of 376 Da was diluted to a stock concentration of 30 mM in PBS.

### Antibodies

Primary antibodies for immunocytochemistry were for claudin-5 (Thermo-Fisher, cat. #34-1600, 1:250), occludin (Thermo-Fisher, cat. #71-1500, 1:250), ZO-1 (Thermo-Fisher, cat. #33-9100, 1:250), CD31 (Abcam, cat. #ab28364, 1:200), GFAP (Covance, cat. #MAB360, 1:200), PDGFRβ (Cell Signalling, cat. #28E1, 1:200), TUJI (Covance, cat. MMS-435P, 1:200) and the GST tag (Proteintech, cat. #66001-2-1g, 1:500). Secondary antibodies were goat-anti-mouse Alexa Fluor 488 (Thermo-Fisher, cat. #A-11029, 1:500), goat-anti-mouse Alexa Fluor 594 (Thermo-Fisher, cat. #A-11005, 1:500), goat-anti-rabbit Alexa Fluor 488 (Thermo-Fisher, cat. #A-1108, 1:500) and goat-anti-rabbit Alexa Fluor 647 (Thermo-Fisher, cat. #A-21245, 1:500). Primary antibodies for western blotting were for claudin-5 (Thermo-Fisher, cat. #34-1600, 1:1,000), occludin (Thermo-Fisher, cat. #71-1500, 1:1,000), ZO-1 (Thermo-Fisher, cat. #33-9100, 1:1,000), GFP (Aves Labs, cat. #GFP-1020, 15,000) and the GST tag (Proteintech, cat. #66001-2-1g, 1:2000). Secondary antibodies were IRDye^®^ 800CW goat-anti-mouse (LiCor, cat. #926-32210, 1:10,000), IRDye^®^ 680LT goat anti-rabbit IgG (H + L) (LiCor, cat. #926-6802, 1:10,000) and horseradish peroxidase (HRP)-conjugated goat anti-chicken IgG (GE Healthcare, cat. #AS09603, 1:10,000).

### Immunocytochemistry

The cells were grown on coverslips and fixed in acetone/methanol (1:1) and pre-incubated in a blocking solution containing 5% goat serum and 1% BSA in PBS. Samples were then incubated with primary antibodies overnight at 4 °C on a rocker. Reactions were visualized by fluorescently labelled anti-mouse or anti-rabbit antibodies. Images were obtained using a 63× objective on a Zeiss LSM 510 META confocal microscope equipped with a 30 mW 405 nm diode laser, a 25 mW 458/488/514 nm argon multiline laser, a 20 mW DPSS 561 nm laser and a 5 mW HeNe 633 nm laser mounted on a Zeiss Axio Observer Z1.

### Western blotting

Confluent cells in T25 flasks were washed twice with PBS, and then lysed with 1× radioimmunoprecipitation assay (RIPA) buffer (Cell Signalling Technologies, cat. #9806) and 200 mM phenylmethylsulfonyl fluoride (Sigma, cat. #P7626) with 1× protease inhibitor cocktail (Roche) for 20 min on ice. Lysates were then homogenized by sonication for 40 s (4× at 20% amplitude, 10 s each) and centrifuged at 10,000 × *g* for 20 min at 4 °C to remove cellular debris before the supernatant was collected for analysis. Total protein content was determined with the Pierce™ bicinchoninic acid protein (BCA) (Thermo-Fisher) assay kit, and samples were prepared by denaturing 15 μg of protein with 8 μl of 5× Laemmli buffer (60 mM Tris/Cl (pH 6.8), 2% SDS, 10% glycerol, 5% ß-mercaptoethanol and 0.01% bromophenol blue) at 95 °C for 5 min. Samples were then loaded onto 4-15% Criterion™ TGX™ Precast Midi Protein Gels (12 + 2 wells, 45 μl) (Bio-Rad) and subjected to SDS-PAGE at 200 V for ~45 min in 10× Tris/Glycine/SDS running buffer (Bio-Rad). Proteins were then transferred onto low-fluorescence 0.45 μm PVDF membranes with transfer buffer (10× Tris/glycine with 10% methanol) (Bio-Rad) for 10 min using the semi-dry Trans-Blot Turbo Transfer System (Bio-Rad). Non-specific binding was blocked with Odyssey^®^ Blocking Buffer (Li-Cor) or 5% skim milk in Tris-buffered saline (TBS) containing 0.1% Tween-20 for 1 h at room temperature followed by incubation with primary antibodies overnight at 4°C. Membranes were then washed 4 times (5 min each) before incubation with fluorescently-labelled secondary antibodies or HRP-conjugated secondary antibodies for 1.5 h at room temperature. Following extensive washes, blots were visualized with an Odyssey Infrared Imager CLX (Li-Cor) and Image Studio™ software (Li-Cor).

### Microbubbles

Biologically inert microbubbles were prepared in-house as described previously [29, 30]. Briefly, 1,2-distearoyl-*sn*-glycero-3-phosphocholine (DSPC) and 1,2-distearoyl-*sn*-glycero-3-phosphoethanolamine-N- [amino (polyethylene glycol)-2000] (DSPE-PEG2000) (Avanti Polar Lipids) were mixed in a 9:1 molar ratio and dissolved in a small amount of chloroform (Sigma) in a glass beaker, followed by evaporation under vacuum (Mivac Quattro) at 22 °C for 20 min. The dried lipid film was then rehydrated in PBS with 10% glycerol to a concentration of 1 mg lipid/ml and heated to 55 °C in a sonicating water bath until fully dissolved. The clear solution was dispensed aseptically into 1.5 ml glass high-performance liquid chromatography (HPLC) vials and the air in each vial was replaced with octafluoropropane (Arcadophta). On the day of the experiment, the HPLC vial was brought to room temperature one hour ahead of the experiment. An equal volume of 0.9% NaCl solution was injected into the vial, followed by agitation in a dental amalgamator for 45 s to generate the MBs. The MBs were characterized for their size and concentration using a Multisizer 4e coulter counter (Beckman Coulter Life Science) as previously described [31].

### FUS application

Cells were cultured on Transwell culture inserts with a pore size of 0.4 μm (Cat. #CLS3470, Corning Inc.) which were placed into a 24-well plate. The plate was then placed on top of a Sonic Concepts H117 ultrasound transducer that was submerged in a water bath adjusted to 24 °C. A proper alignment of the focus was performed. 20 μl of microbubbles were added to the top chamber of each well and exposed to FUS (286 kHz centre frequency, 50 cycles/burst, burst period 20 ms and a 120 s sonication time). A range of pressures were tested and the input power was based on calibration of the system with a calibrated needle hydrophone (NPL, UK). For GST-cCPEm treatment, cells were incubated in a 37 °C incubator prior to FUS^+MB^ treatment.

### Transendothelial electrical resistance and paracellular permeability

TEER across cellular monolayers was measured with chopstick electrodes in 24-well Transwell inserts using the Millicell^®^ ERS-2 Voltohmmeter (Merck Millipore). The absolute TEER of eGFP-hCldn5-MDCK II cells before treatment was recorded as a baseline reading. Resistance of the blank filter was subtracted and then multiplied by the surface area of the membrane for calculation of the final TEER values. All experiments were carried out in triplicates, and data are expressed as mean ± SEM of two independent experiments. The permeability to fluorescently labelled tracer molecules was also used to estimate the paracellular opening in monolayers due to treatments. 10 μM NaFl or 0.5 mg/ml FD4 or 0.5 mg/ml FD40 was added to the apical chambers and incubated at 37 °C. After 2 h, 100 μl samples from the bottom chambers were collected for reading in a fluorescence plate reader (CLARIOstar-BMG Labtech) at 483 ± 14 nm excitation and 530 ± 30 nm emission wavelengths.

### Viability assays

To determine cell viability post-treatment, a cytotoxicity assay (CellTiter 96, Promega) was performed following the manufacturer’s instructions. For all experimental conditions, premixed MTT (3-(4,5-dimethylthiazol-2-yl-2,5-diphenyltetrazoliumbromide; 0.25 mg/ml, Thermo-Fisher, cat. # M6494) solution was added to the wells, followed by incubation for 50 min. DMSO was then added to the culture wells, and the plate was incubated at RT on a rocker for 10 min. Absorbance was recorded at 570 nm using a plate reader (CLARIOstar-BMG Labtech). Values were measured as fold change compared to the untreated samples and plotted as mean ± SEM.

### Statistical analysis

Statistical analysis was performed with GraphPad Prism 9 using an unpaired student’s t-test or two-way ANOVA with Sidak’s or Turkey’s multiple comparison test. All data presented as mean ± SEM. Values outside two standard deviations from the mean were excluded.

### Ethics

Animal experimentation (microvessel isolation) was approved by the Animal Ethics Committee of the University of Queensland (approval numbers QBI/348/17/NHMRC and QBI/554/17/NHMRC). The animals were housed in specific pathogen-free cages and maintained on a 12 h light/dark cycle, with unlimited access to food and water.

## Results

### Establishment and characterization of eGFP-hCldn5 MDCK II cells

To manipulate BBB opening *in vitro*, we first evaluated a range of cell lines which are routinely used for these purposes. Of these, human iBECs are a suitable i*n vitro* system because they form claudin-5-containing TJs, display a cobblestone-like morphology and have a high TEER (in the order of 4,000 Ω·cm^2^) [15], but the challenge is to ensure a constant supply of these cells. MDCK II cells, on the other hand, have been reported to display a much lower TEER of up to 200 Ω·cm^2^ [32–34]. They express TJ proteins such as occludin and ZO-1, as well as several claudins including claudin-1, claudin-2 and claudin-4, but not claudin-5 [35]. Using a lentiviral approach, we generated a stable eGFP-hCldn5-MDCK II cell line **(Fig. 1A,B)**, establishing eGFP-expressing cells as a negative control. Both cell lines [34] displayed a cobblestone-like appearance, but whereas eGFP was localized to the cytoplasm in eGFP-MDCK II cells, the fusion protein was localized to the membrane in eGFP-hCldn5 MDCK II cells, as expected of a TJ protein **(Fig. B-D)**. Importantly, claudin-5 did not localize to the plasma membrane when the cells were not in contact (i.e., when no TJs formed) **(Fig. 1C)**. Claudin-5 could also be visualized with an anti-claudin-5 antibody, and the protein was found to be colocalized at cell-cell borders with both occludin and ZO-1, indicative of the protein being a TJ component **(Fig. 1E-G)**.

As a further validation, we performed a western blot analysis of eGFP-hCldn5-MDCKII, eGFP-MDCKII and untransfected cells, using antibodies for claudin-5, eGFP, occludin and ZO-1 **(Fig. 2A, Suppl. Fig. 3)**. The cells expressed occludin and ZO-1. With eGFP having a molecular weight of 27 kDa and claudin-5 of 17 kDa, this revealed a claudin-5 fusion protein of an apparent molecular weight of 44 kDa.

**Figure 2:**
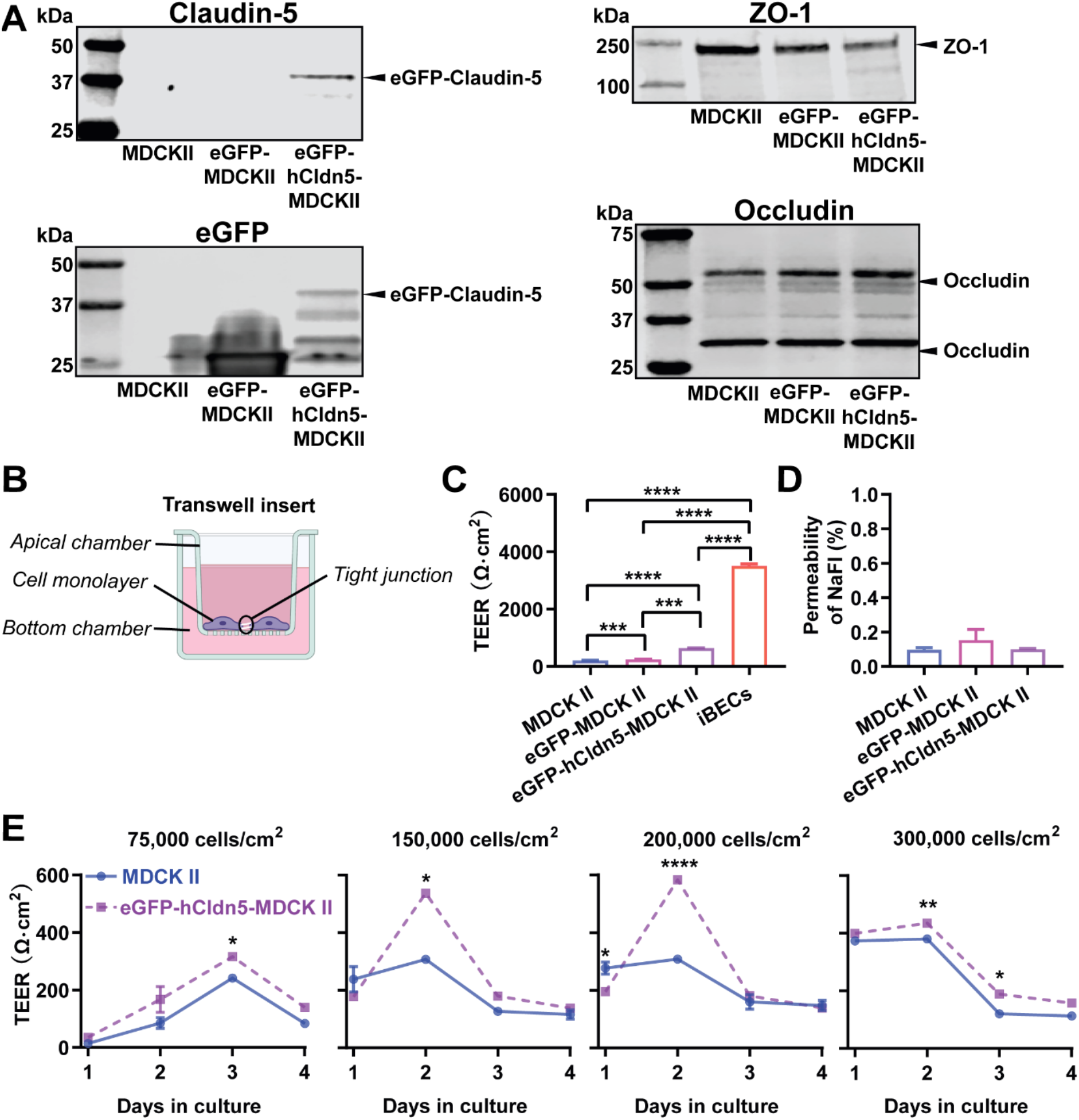
eGFP-hCldn5-MDCK II cells exhibit a tight monolayer as determined by TEER and cargo leakage. **(A)** Western blotting with either a claudin-5 or GFP antibody reveals expression of the eGFP-hClaudin5 fusion protein in eGFP-hCldn5-MDCK II but not eGFP-MDCK II cells. Expression of the tight junction proteins ZO-1 and occludin is also shown. **(B)** Scheme of Transwell insert to measure TEER. eGFP-hCldn5-MDCK II cells display a four-fold higher TEER than MDCK II cells. iBEC cells are included for comparison. **(C)** All three MDCK II cell lines show a < 0.2% permeability for sodium fluorescein (NaFl), indicating a tight BBB. **(D)** TEER of eGFP-hCldn5-MDCK II as both a function of cell density and days in culture. TEER values are shown as Ω·cm^2^ and results are expressed as mean ± SEM. N= 8-14 per condition. Statistical significance was determined as unpaired student’s t-test (*p<0.05, **p<0.01, ***p<0.001 and ****P<0.0001).

Next, we determined the TEER of eGFP-hCldn5-MDCK II cells compared to the parental MDCK II cell line and iBECs, culturing the cells on Transwell inserts. The TEER of iBECs was measured to be 3450 Ω·cm^2^, consistent with what has been previously reported [15]. Importantly, the mean TEER of eGFP-hCldn5-MDCK II cells was 636 ± 40.5 Ω·cm^2^, i.e., 3-fold higher than that of MDCK II cells (193 Ω·cm^2^) **(Fig. 2B)**. Of note, the TEER of eGFP-MDCK II control cells was slightly higher than for untransfected cells, possibly reflecting the fact that eGFP-MDCK II tended to overgrow, which was neither observed for the parental MDCK II cells nor for eGFP-hCldn5-MDCK II cells. The permeability characteristics of the eGFP-hCldn5-MDCK II and MDCK II cells were also determined by measuring the paracellular permeability of the cells grown on Transwell inserts to NaFl and fluorescently labelled 4 kDa dextran (FD4). This analysis revealed that less than 0.2% of NaFl (MW 376 Da) passed to the lower chamber of the Transwell insert after 2 h, indicating a high level of barrier integrity **(Fig. 2C)**. No leakage was seen for FD4 (MW 4 kDa) across the cellular monolayer **(data not shown)**. We also found that the TEER value was a function of cell density (testing four seeding densities ranging from 75,000 to 300,000 cells/cm^2^) and days in culture in that the highest TEER was obtained at 2 days in culture for cells seeded at 150,000 or 200,000 cells/cm^2^ **(Fig. 2D)**. We thereby established a model that allowed us to explore barrier function by measuring the TEER and permeabilities of the monolayer.

### cCPEm and mC5C2 display a different time course of barrier opening in eGFP-hCldn5 MDCKII cells

The modified toxin GST-cCPEm has been reported to bind to ECL2 of claudin-5 with nanomolar affinity, whereas the peptide mC5C2 binds to ECL1 with a suggested nanomolar affinity **(Fig. 3A)** [17]. We synthesized cCPEm and generated a fusion protein with cCPEm being carboxy-terminally fused to GST. As we did not remove the GST moiety, this allowed us to visualize binding to the TJs using an anti-GST antibody. By isolating murine cerebral microvessels **(Suppl. Fig. 4A-D)** we had a specimen in which we were able to observe BBB binding of GST-cCPEm, which colocalized with mCld-5 **(Suppl. Fig. 4E)**. mC5C2 was unlabelled so its binding could not be visualized. To determine how cCPEm and mC5C2 weaken the BBB, we tested the effects of different concentrations of the binders in time-course experiments in eGFP-hCldn5-MDCK II cells grown to confluency in 6.5-mm Transwell insets. A viability test proved that the two binders did not cause overt cytotoxicity **(Suppl. Fig. 5)**.

**Figure 3:**
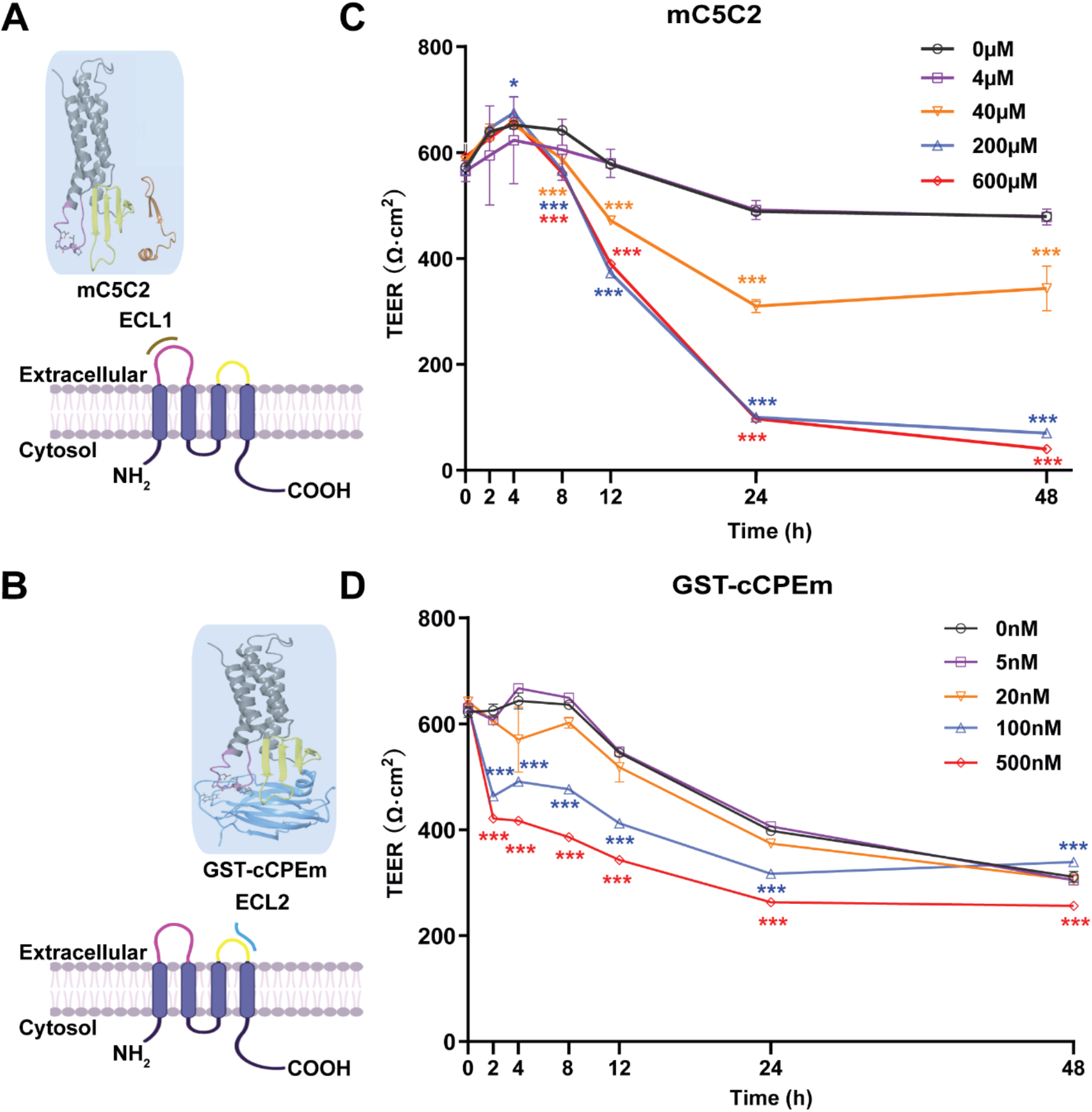
Incubation with mC5C2 and GST-cCPEm reveals differences in the reduction of the relative TEER in eGFP-hCldn5-MDCK II cells. **(A,B)** Schematic diagram showing mC5C2 binding of the extracellular loop 1 (ECL1) and cCPEm binding ECL2 of claudin-5. The homology model of claudin-5 was created in Swiss-Model using human claudin-9 (PDB ID 6OV2) as template. The cCPEm structure was extracted from the same PDB entry (6OV2). mC5C2 was placed in proximity to the model of claudin-5 whereas cCPEm was docked to claudin-5 for schematic purposes only. The molecular structures were generated using Maestro (Schrödinger Release 2020-4, New York, 2020). **(C,D)** Incubation of eGFP-hCldn5-MDCK II with mC5C2 and GST-cCPEm causes concentration- and incubation-time-dependent reductions in the relative TEER. N=6 of each condition. Two-way ANOVA with Sidak’s multiple comparison test (*p<0.05, and ***p<0.001).

Cells were consistently plated at a density of 200,000 cells/cm^2^ and treated 2 days after plating. We tested mC5C2 at 0, 4, 40, 200 and 600 μM, and GST-cCPEm at 0, 4, 20, 100 and 500 nM concentrations, measuring both absolute TEER (in Ω·cm^2^) at 0, 2, 4, 8, 12, 24 and 48 h of incubation **(Fig. 3 B,D)** and relative TEER **(Suppl. Fig. 6)**. Compared with the peptide-free medium control, the TEER of eGFP-hCldn5-MDCK II cells displayed a time- and concentration-dependent decrease following treatment with mC5C2 and GST-cCPEm **(Fig. 3B,D)**. We observed a weakening of the BBB as measured by TEER for the mC5C2 peptide after an 8 h incubation, and for GST-cCPEm after 4 h, with some indication of a drop at the higher concentrations already after 2 h. Together, there was a clear concentration-dependent effect for both treatments. Of note, the drop in TEER was more pronounced for the peptide with no indication of BBB closure even after 48 h, whereas for GST-cCPEm, the drop in TEER was less pronounced.

### Focused ultrasound treatment with acoustic pressures of 0.3 and 0.4 MPa achieve reliable barrier opening

We next explored FUS in the presence of microbubbles (FUS^+MB^) which is known to open the BBB in a time scale of seconds [36, 37] **(Fig. 4A)**. The average diameter of MBs used in our study was 1.060 ± 0.64 μm, with a concentration of ~ 9.0 × 10^9^ MBs/ml **(Suppl. Fig. 7A)**. For iBEC cells, we had previously shown cells detach from the monolayer at pressures of 0.3 MPa, and that at 0.15 MPa, barrier opening is achieved without destroying the monolayer and creating larger holes [15]. MDCK II cells are more robust, which is why we tested a range of slightly higher pressures (up to 0.4 MPa). We found that at 0.3 and 0.4 MPa, the barrier was opened as demonstrated by a reduction in absolute TEER **(Fig. 4B),** without causing damage to the cell layer as demonstrated via an MTT assay for all conditions **(Suppl. Fig. 5)**. To better compare the FUS^+MB^ data with those obtained for GST-cCPEm and mC5C2, we next used the same time points as above (0, 2, 4, 8, 12, and 24 h) to better understand barrier opening and closure. By determining the absolute **(Fig. 4C)** and relative changes to TEER **(Suppl. Fig. 7B)** this revealed that the barrier closed within 12 h for the conditions tested.

**Figure 4:**
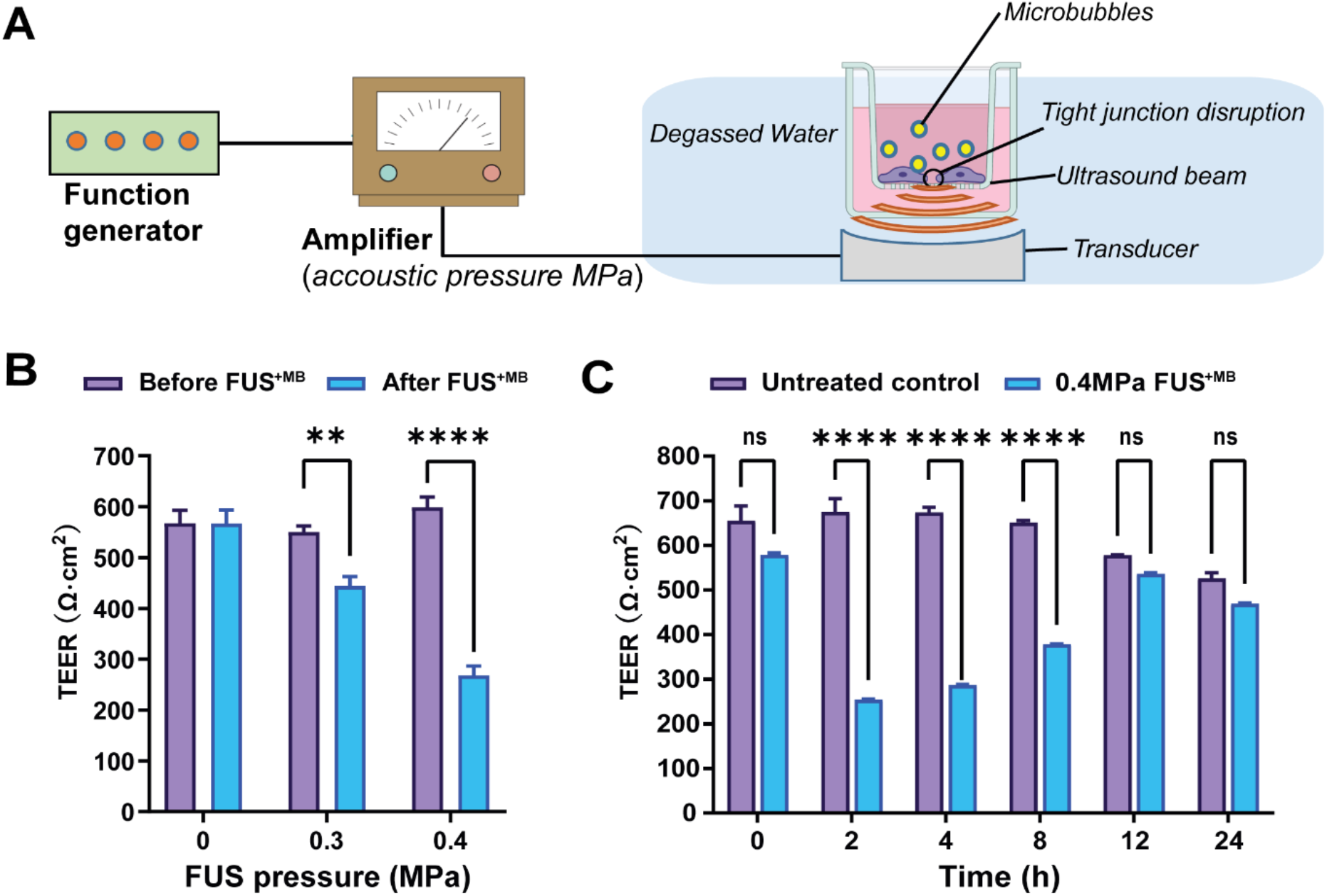
Focused ultrasound with microbubbles (FUS^+MB^) leads to a rapid opening of the barrier followed by closure within 12 hours. **(A)** Schematic diagram of how ultrasound is delivered to the cells. **(B)** TEER measurement as a function of acoustic pressure (in MPa). **(C)** Absolute TEER measurement as a function of incubation time shown for the 0.4 MPa condition. N=3-6 for each condition. Two-way ANOVA with Sidak’s multiple comparisons tests (**p<0.01 and ****P<0.0001).

### Preincubation with the BBB binder cCPEm enhances focused ultrasound-mediated barrier opening

We next examined whether pre-treatment with a BBB binder would lower the threshold required for FUS^+MB^ to achieve barrier opening and achieve a greater degree of opening at a pressure which results in weak opening in the absence of GST-cCPEm. Several parameters were explored to narrow down suitable conditions **(data not shown)**. Given that GST-cCPEm, at the concentrations used, achieved barrier opening faster than mC5C2 as measured by TEER, we pre-incubated the eGFP-hCldn5-MDCK II cells with 100 nM GST-cCPEm for 2 h, followed by FUS^+MB^ at 0.1, 0.2 and 0.3 MPa. GST-cCPEm and FUS^+MB^ only conditions were included as controls **(Fig. 5A)**. TEER was determined at the start of the experiment (0 h, +/− GST-cCPEm) and after 2h (+/− sonication). Leakage of 40 kDa FITC-labelled dextran FD40 was also determined after 2h. We found that for all conditions, TEER was reduced compared to the untreated control.

**Figure 5:**
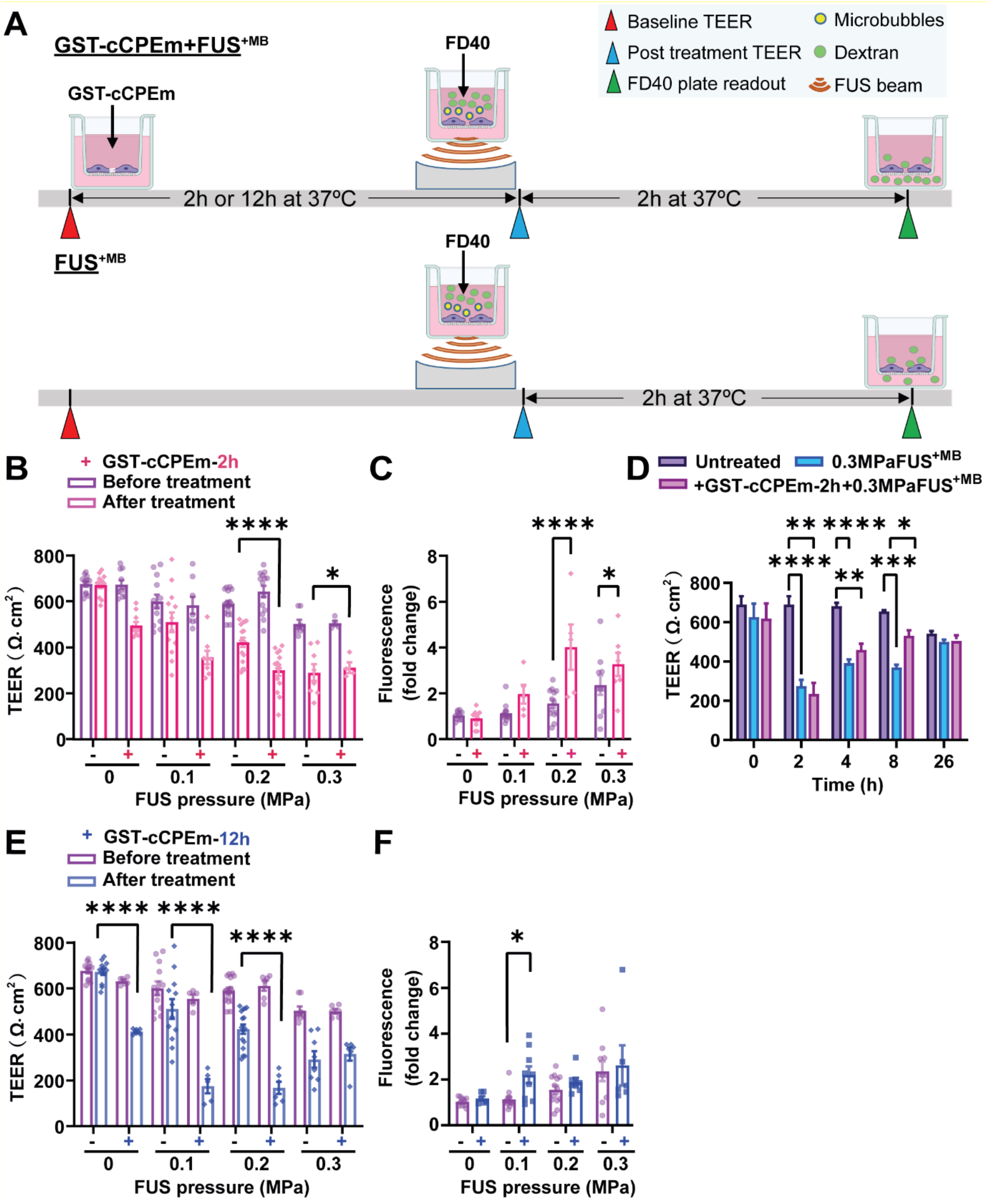
Preincubation with GST-cCPEm lowers the acoustic pressure required for focused ultrasound-mediated barrier opening. **(A)** Experimental scheme. **(B)** Absolute TEER measurement and **(C)** FD40 leakage of eGFP-hCldn5-MDCK II cells preincubated with 100 nM GST-cCPEm for 2 h, followed by FUS^+MB^ at the indicated pressures. **(D)** TEER measurement to assess barrier closure when GST-cCPEm was removed followed treatment. **(E)** Absolute TEER measurement and **(F)** assessment of FD40 leakage of eGFP-hCldn5-MDCK II cells preincubated with 100 nM GST-cCPEm for 12 h, followed by FUS^+MB^ at the indicated pressures. N≥5 from at least two independent experiments. Two-way ANOVA with Tukey’s and Sidak’s multiple comparisons tests (*p<0.05, **p<0.01, ***p<0.001 and ****P<0.0001).

At 0.3 MPa, GST-cCPEm had no impact on the extent of TEER reduction; however, by progressively dropping the acoustic pressure to 0.2 and then 0.1 MPa, the TEER reductions became more pronounced, with an additional 43.7% and 37.7% TEER reduction after preincubation with GST-cCPEm at the 2 h timepoint. A two-way ANOVA revealed a significant effect of FUS^+MB^ F(3,75)=45.29, p<0.0001 and a significant effect of GST-cCPEm on TEER F(1,75)=43.47, p<0.0001 **(Fig. 5B)**. The changes in TEER were reflected by changes in permeability as determined for FD40 at 2 h post FUS^+MB^, which was increased by 158.7% and 73.6% when pre-incubated with GST-cCPEm compared with FUS^+MB^ alone at 0.2 MPa and 0.1 MPa, respectively. A two-way (FUS^+MB^ pressure) x (GST-cCPEm) ANOVA on FD40 leakage found there was a significant main effect of FUS^+MB^ (F_3,63_=17.49, p<0.0001). There was a significant main effect of GST-cCPEm on FD40 leakage (F_1,63_=23.72, p<0.0001) indicating GST-cCPEm increases the permeability of the monolayer to FD40 separately to the effect of FUS^+MB^. The interaction was significant (F_3,69_=5.74, p=0.0015) indicating that the effect of GST-cCPEm is different depending on the ultrasound pressure applied with leakage being enhanced by GST-cCPEm most successfully at 0.2 MPa FUS^+MB^ (p<0.0001) and also at 0.3 MPa FUS^+MB^ (p<0.03 Sidak multiple comparison’s test) **(Fig. 5C)**. Importantly, to explore whether the barrier integrity of treated monolayers could be restored, we removed the GST-cCPEm from the cell medium after the FUS treatment and monitored TEER for 24 h. No significant difference in absolute TEER values was observed between treated groups and the untreated control after 24 h **(Fig. 5D).** The cell monolayer integrity was restored with TEER reaching around 85% of the baseline value after 24 h.

We then extended our study and prolonged the incubation time for the toxin to 12 h. Our results revealed that at a pressure of 0.1 and 0.2 MPa, when eGFP-Cldn5-MDCK II cell monolayers were preincubated with GST-cCPEm for 12 h, the extent of TEER reduction for the combination treatment was a further 166.5% and 160.6%, compared with cells treated with 0.1 and 0.2 MPa FUS^+MB^, respectively. A two-way (FUS^+MB^ pressure) x (GST-cCPEm) ANOVA on the TEER recorded at 12h post FUS^+MB^ found there was a significant main effect of FUS^+MB^ such that there was a decrease in TEER with increasing pressure (F_3,63_ = 23.07, p<0.0001). There was a significant main effect of GST-cCPEm incubation revealing that preincubation with GST-ccCPEm lowers the TEER separately to the effect of FUS^+MB^ (F_1,63_=73.61, p<0.0001). There was a significant FUS^+MB^ x GST-cCPEm interaction such that GST-cCPEm had a greater effect on TEER at low ultrasound pressures which was confirmed by Sidak multiple comparison’s test which found that for each pressure the TEER was significantly lower when the cells were treated with combined FUS^+MB^ and GST-cCPEm indicating our hypothesis that GST-cCPEm would increase the amount of BBB opening at lower pressures of 0.1 MPa and above **(Fig. 5E)**. The changes in TEER were reflected by changes in permeability in that the leakage of FD40 was increased by FUS^+MB^ (p=0.0017, two-way ANOVA) and GST-cCPEm (p=0.046, two-way ANOVA) **(Fig. 5F).**

Together, these two time-course experiments reveal that pre-incubation with a BBB binder lowers the threshold for the acoustic pressure required to open TJs, suggesting that pre-treatment with a claudin-5 binder followed by FUS may achieve safe BBB opening *in vivo*.

## Discussion

Focused ultrasound with microbubbles (FUS^+MB^) is an emerging technology for the treatment of Alzheimer’s disease and other neurological conditions [6]. Therapeutic ultrasound, without delivering therapeutic agents, has been shown to reduce the amyloid-β and tau pathology of Alzheimer’s disease and improve memory and motor functions [29, 38, 39]. Currently, this technology is being explored in a range of clinical trials in patients with Alzheimer’s disease. The methodology is used in combination with intravenously injected MBs, which respond to ultrasound by going through repeated cycles of expansion and contraction [4]. This causes the severing of TJs between adjacent brain endothelial cells, thereby achieving transient BBB opening. However, given that the useful therapeutic pressure range is only within a narrow window, there is the inherent risk of brain damage due to the extravasation of blood cells in response to high pressures [40–42]. When the pressure is too low, there is no functional BBB opening and no cargoes are taken up, and as the pressure increases, cargoes can be taken up increases, until due to the interaction with microbubbles a pressure threshold is reached that causes inertial cavitation of the microbubbles with ensuing damage [43]. Agents which have been safely delivered into the brain parenchyma by FUS include dextrans of various sizes [44, 45], chemotherapeutic drugs such as doxorubicin or methotrexate [46, 47], antibodies in various formats [48–51], and as viral vectors for gene therapy [52]. Here we asked whether it would be possible to increase permeability at a given FUS pressure when combined with compounds targeting TJ proteins such as claudin-5. In a clinical setting this would offer the possibility to use lower acoustic pressures to achieve the same uptake.

To investigate this, we first established an *in vitro* hCldn5-MDCK II cell culture system in which we explored two binders of claudin-5, mC5C2 and GST-cCPEm, both of which weaken the BBB over time. Compared with FUS, the two binders showed a different profile for the conditions we tested. Whereas FUS caused rapid BBB opening followed by recovery after 12 h within the tested pressure range (0.1-0.4 MPa), GST-cCPEm achieved opening as measured by a change in TEER only after 2-4 h, and mC5C2 needed even longer (up to 8 h) to open the barrier within the examined concentration range. No closure was detected for mC5C2 within a time window up to 48 h after treatment. Whether for GST-cCPEm there was a slight indication of recovery at around 48 h is difficult to conclude given that there a marked baseline reduction in TEER between 36 and 48 h because the culture medium is not being changed and cell health deteriorates after this incubation time **(Fig. 3D**). Weakening of the barrier was assessed by measuring the TEER and transepithelial passage of fluorescent tracers. Although both are indicators of TJ integrity, they reflect different experimental parameters. The TEER reflects the ionic conductance of the paracellular pathway, whereas the flux of fluorescent tracers indicates paracellular water flow and the pore size of the TJs [12]. A limitation of our study is that whereas FUS is given as a short pulse with a duration of 2 min, incubation with either mC5C2 or GST-cCPEm was for the entire duration of the experiment. In an *in vivo* setting, the question arises how long GST-cCPEm would persist in the circulation. It seems therefore to be more relevant to determine e.g. in mice (where the compound cannot be washed away) to determine at which concentrations and how much before the FUS^+MB^ treatment GST-cCPEm has to be provided to affect BBB opening.

Our approach was guided by FDA guidelines which require a TEER threshold of at least 100 Ω·cm^2^ for any cell line used as a cell permeability tool [53]. By generating an MDCK II cell line that expresses fluorescently labelled claudin-5, we were demonstrating an optimally high TEER two days post-seeding, whereas Caco-2 cells, for example, require more than two weeks [54]. We expressed human claudin-5 in our cell line of canine origin as no canine claudin-5 has been reported. Another consideration was that, in a clinical setting, TJs express the human form of the protein. Murine claudin-5 has been used for the transfection of human brain endothelial (hCMEC/D3) cells, demonstrating integrity of the TJ strand network [55]. Human and murine claudin-5 are highly homologous (98.2%) and their two ECLs show 100% homology between species [56]. Therefore, we would not expect any functional difference between human claudin-5 and murine claudin-5 transduced MDCK II cells with respect to barrier formation. An obvious limitation of our cellular system is that it lacks other cells that are part of the NVU such as astrocytes and pericytes; however, future studies will determine the extent to which the findings obtained in a simple MDCK II cell system can be translated *in vivo*.

In an experimental or clinical *in vivo* setting, we foresee that the approach discussed here may be more generally applicable. Any peptide or small molecule compound that not only binds to but also weakens the interaction of TJ molecules such as claudin-5, or even adherens junction proteins, may be useful for lowering the threshold at which therapeutic ultrasound can be safely delivered. Alternative strategies could involve the combination of FUS with bradykinin (which increases the number of pinocytotic vesicles and hence facilitates transcytoplasmic transport), vasodilators such as papaverine (which reduce transcription and hence, the translation of proteins, including those that constitute TJs), or endothelial monocyte-activating polypeptide II (EMAP-II, which exerts a similar effect to papaverine) [57–59]. In our study we have not addressed transcytoplasmic transport. With regards to the effects on the transcription and translation of TJ proteins such as claudin-5, the effects observed in the aforementioned studies are the consequence of a signal transduction cascade which presumably has multiple effects. Our approach is different in that it targets TJ proteins directly. Here, the BBB binders destabilize the BBB by forming a wedge **(Fig. 6B)**, lowering the requirements for ultrasound to open the barrier, which ultimately should allow for the more effective and safer uptake of therapeutic agents by the brain.

**Figure 6:**
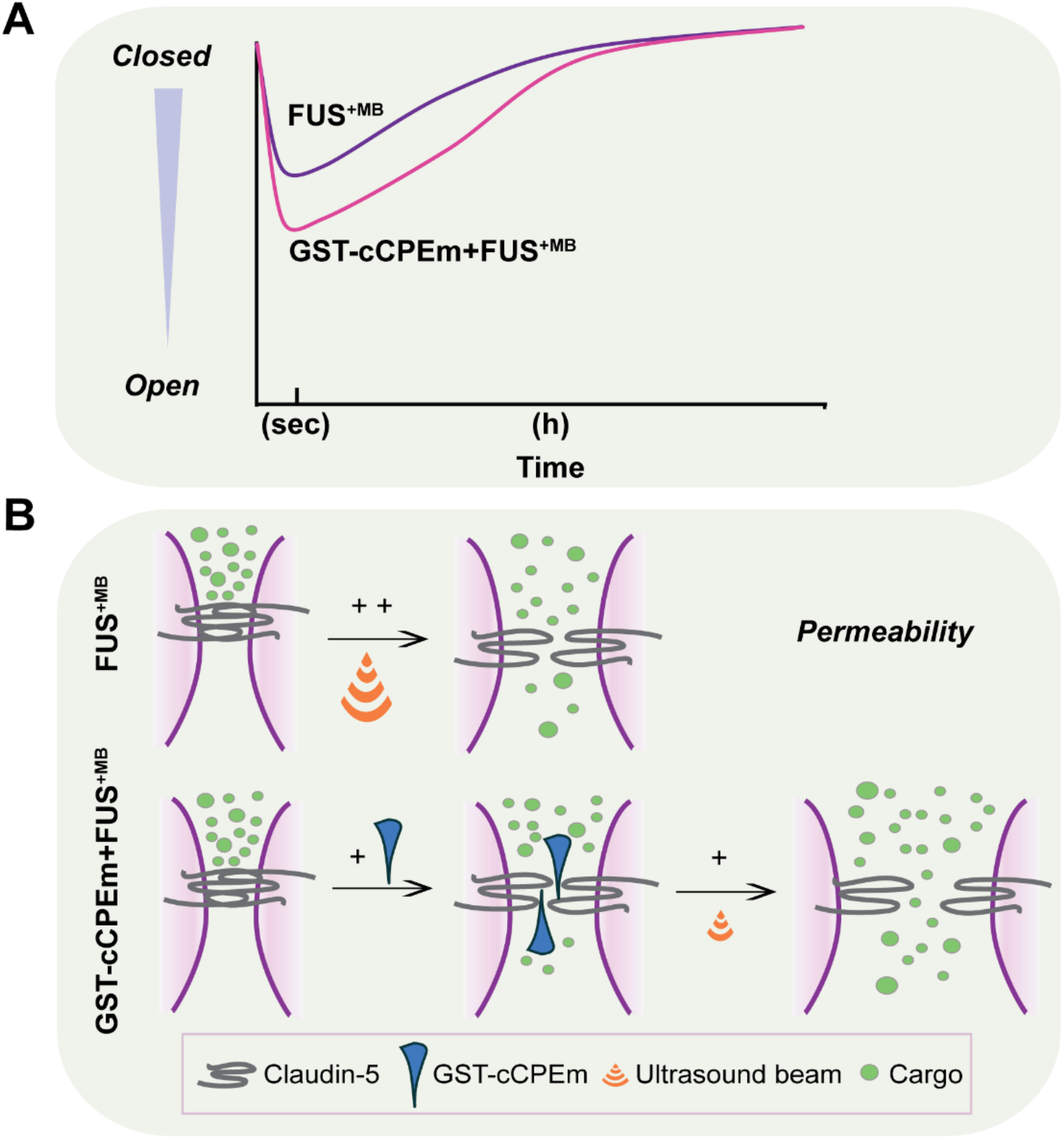
Summary scheme of combinatorial treatment. **(A)** Scheme of opening effect of cells pre-treated with or without GST-cCPEm followed by FUS^+MB^. **(B)** GST-cCPEm forms a wedge which destabilizes claudin-5 and thereby the TJ, making it easier for the ultrasound pressure wave to open the barrier, with potential implications for FUS-mediated drug delivery.

## Abbreviations

BBB: blood brain barrier
FUS: focused ultrasound
MBs: microbubbles
TJs: tight junctions
TEER: transendothelial electrical resistance
CNS: enctral nervous system
NVU: neurovascular unit
BCECs: brain capillary endothelial cells
JAM: adhesion molecule
ESAMs: endothelial cell adhesion molecules
TM: transmembrane domains
ECL: extracellular loops
hiPSCs: human induced pluripotent stem cells
iBECs: human induced pluripotent stem cells derived endothelial cells
hCMEC/D.3: human cerebral microvascular endothelial cells
MDCK II: Madin-Darby Canine Kidney
FUS+MB: FUS, in conjunction with biologically inert gas-filled microbubbles
ECM: endothelial cell medium
GFAP: glial fibrillary acidic protein
PDGFRβ: platelet derived growth factor receptor beta
TUJ1: class III beta-tubulin
FD: Fluorescein isothiocyanate-dextran
NaFl: sodium fluorescein
DSPC: 1,2-distearoyl-sn-glycero-3-phosphocholine
DSPE-PEG2000: 1,2-distearoyl-sn-glycero-3-phosphoethanolamine-N-[amino (polyethylene glycol)-2000]
HPLC: high-performance liquid chromatography
RT: room temperature
MTT: 3-(4,5-dimethylthiazol-2-yl-2,5-diphenyltetrazoliumbromide

## Credit author statement

LC, RS, JL and JG designed the experiments; LC, RS, JL, BA and TP performed the experiments; LC, RS, JL analyzed the data; LC and JG wrote the manuscript, with editorial input from all authors.

## Acknowledgements

We thank Adam Briner for advice with cloning, Rowan Tweedale and Dr. Andrew Kneynsberg for critical reading of the manuscript. We would also like to thank the QBI Microscopy facility for assistance with imaging.

## Graphical abstract

Preincubation of a BBB cell line with a claudin-5 binder (GST-cCPEm) acted like a wedge for ultrasound in opening the barrier, with potential *in vivo* implications for drug delivery to the brain.

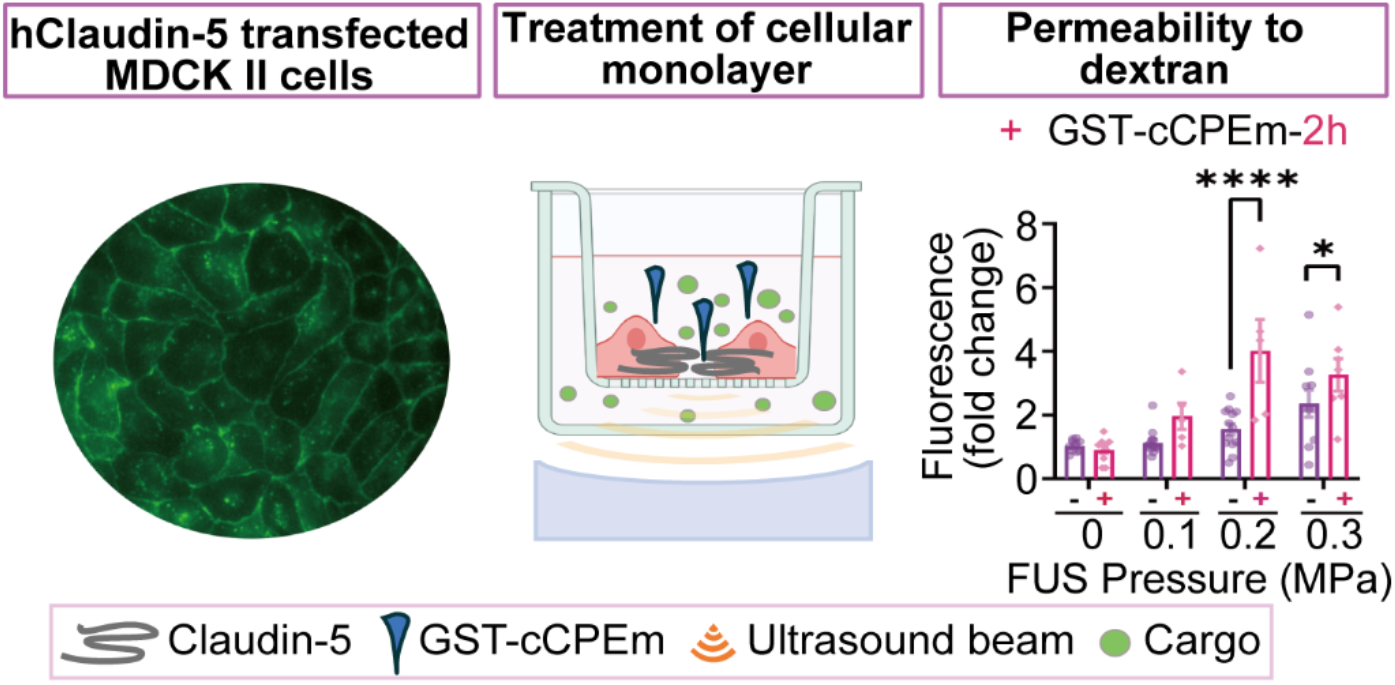

